# Azacytidine targeting SARS-CoV-2 viral RNA as a potential treatment for COVID-19

**DOI:** 10.1101/2021.09.01.458475

**Authors:** Xian Lin, Xianliang Ke, Lin Xia, Tianying Zhang, Hualong Xiong, Binghai Zhao, Wen Liu, Quanjiao Chen, Chong Tang

**Affiliations:** CAS Key Laboratory of Special Pathogens and Biosafety, Wuhan Institute of Virology, Center for Biosafety Mega-Science, CASCIRE, Chinese Academy of Sciences, 430071, Wuhan, China; Department of Pathophysiology, School of Medicine, Beihua University, Jilin 132011, Jilin, China; Fujian Provincial Key Laboratory of Innovative Drug Target Research, School of Pharmaceutical Sciences, Xiamen University, Xiamen, Fujian 361102, China; State Key Laboratory of Molecular Vaccinology and Molecular Diagnostics, National Institute of Diagnostics and Vaccine Development in Infectious Diseases, School of Public Health, Xiamen University, Xiamen, Fujian 361102, China

## Abstract

The COVID-19 pandemic is a global health disaster. Moreover, emerging mutated virus strains present an even greater challenge for existing vaccines and medications. One possible solution is to design drugs based on the properties of virus epigenome, which are more common among coronaviruses.

Here, we reported an FDA-approved drug for myelodysplastic syndrome, azacytidine (5Aza), limited virus infection and protected mice against SARS-CoV-2. We demonstrated that this antiviral effect is related to 5Aza incorporation into viral RNA, which disrupt m5C RNA methylation modification profile. This work suggests that targeting viral epigenomes is a viable therapeutic strategy, potentially opening new pathways for treating COVID-19.

---

As an FDA-approved drug for myelodysplastic syndrome, 5Aza is a structural analog of cytidine (Fig.S1). It exhibits antiviral effects against several viruses. However, it is unknown whether this extends to SARS-CoV-2.

Here, we demonstrated that 5Aza shows potent antiviral activity in Vero E6 cells, with half-maximal inhibitory concentration (IC_50_) = 6.99 μM and selective index (SI) = 20.41 (Fig. 1a). The viral titer in cell supernatants detection (Fig.1b) and indirect immunofluorescence assay (IFA) against viral N protein (Fig.1c) revealed 5Aza restricted SARS-CoV-2 infection in a dose-dependent manner. Time-of-drug-addition assay (Fig.S2a) showed that 5Aza functioned after virus entry (Fig.S2b, c, and d). The virus replication is still 50% suppressed when 5Aza at 32 μM is added 24 hours after infection (Fig. S3).

**Fig. 1.**
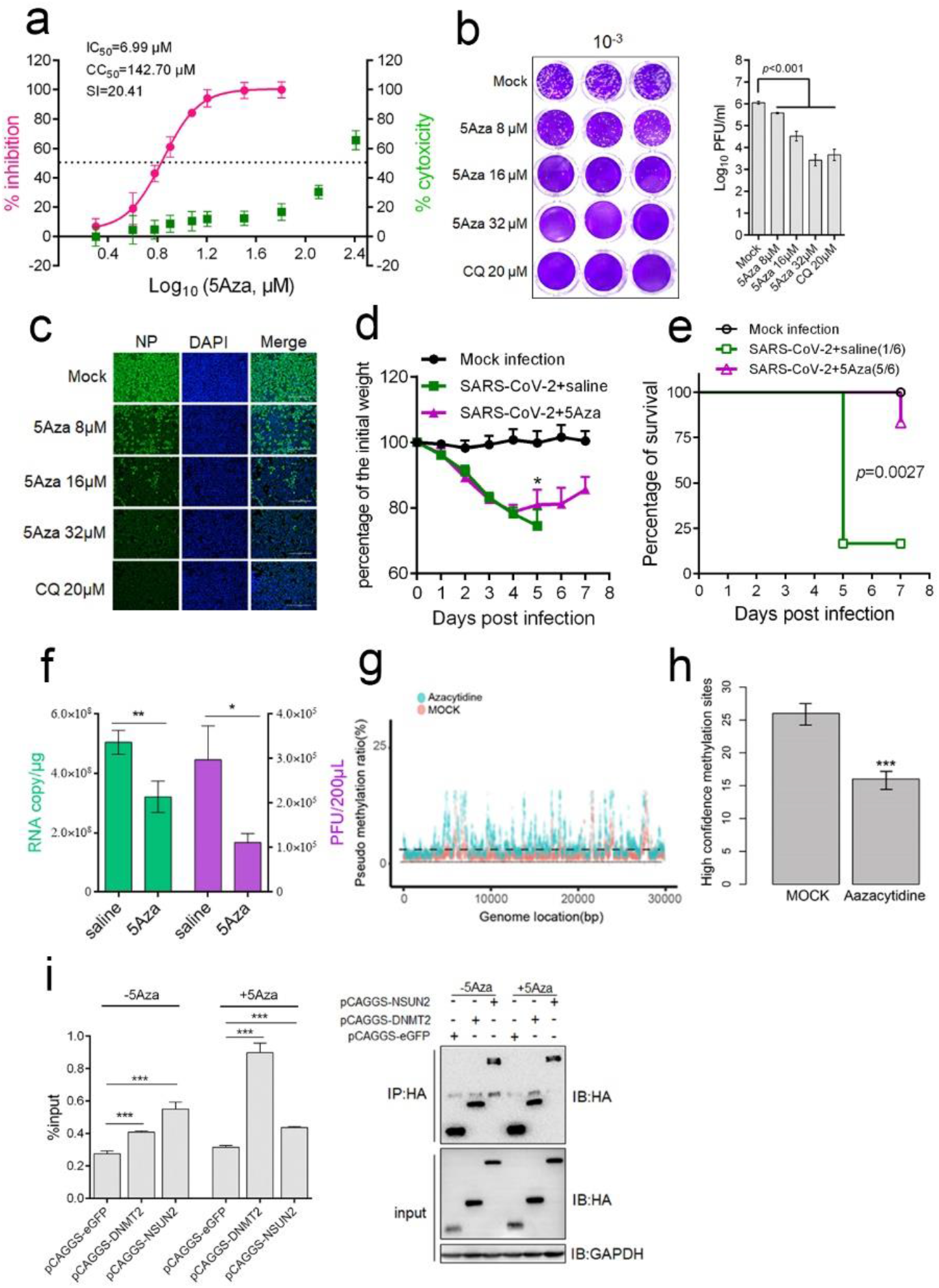
Azacytidine targets SARS-CoV-2 RNA to inhibit virus infection. **a, b, and c** The anti-SARS-CoV-2 effect of Azacytidine *in vitro*. Vero E6 cells were infected with SARS-CoV-2 at MOI = 0.2 in the presence of different doses of Azacytidine. The chloroquine was used as a drug control. At 24 hpi, cell supernatants were collected. **a** IC_50_ and EC_50_ were calculated by detecting viral RNA through RT-PCR and CCK-8 assay, respectively. Left and right Y-axes represent mean % inhibition of virus and % cytotoxicity of 5Aza, respectively. **b** Cell supernatants were collected at 24 hpi, and the viral titers were measured using the plaque assay. **c** Immnofluorescence microscopy of virus infection via probing N protein of SARS-CoV-2. Bars, 200 μM. **d, e,** and **f** The anti-SARS-CoV-2 effect of Azacytidine *in vivo*. 6–7 weeks old female BALB/c mice were randomly divided into three groups with 9 mice per group. Mice were intranasally challenged with 2 × 10^3^ PFU MA-SARS-CoV-2 in 50 μl DMEM (infection groups) or equal DMEM (mock infection). At 1 dpi, mice were intraperitoneally injected with 2 mg/kg 5Aza (SARS-CoV-2+5Aza group) or an equivalent volume of sterile saline (SARS-CoV-2+saline and mock infection groups), once daily for seven consecutive doses. **d** body weight was measured daily for 7 d (n=6), mice with 25% body weight loss were considered moribund and euthanized. Note: the remaining one mouse (body weight loss less than 25%) in SARS-CoV-2+saline group was not recorded at dpi 6 and 7. **e** the survival rates of mice (n=6); mice with more than 25% of body weight loss in **d** were considered moribund and euthanized. **f** at 4 dpi, three mice per group were euthanized for detecting viral RNA copy and virus titer in the lungs. **g** The pseudo m^5^C locations indicated incorporated azacytidine. Vero-E6 cells were infected with 1 moi SARS-CoV-2 in the presence of 10 μM 5Aza or saline. After 12 h, total RNA was isolated and comparison of 5Aza-treated viral RNA with the control using bisulfite sequencing was performed; the non-overlapping points are pseudo m^5^C locations that indicate where 5Aza was incorporated. **h** RNAs of cells infected with SARS-CoV-2 that treated with or without 5Aza were subjected to Nanopore direct RNA sequencing. **i** Vero E6 cells transiently overexpressing DNMT2, NSUN2, or GFP with a HA tag were infected with 0.2 moi SARS-CoV-2 for 20 h, in the presence of 16 μM 5Aza or not. Lysates were prepared and split for incubation with mouse anti-HA antibody. Co-precipitated RNA was analyzed by qRT-PCR using primer sets targeting viral S gene. The level of viral RNA amplicon was determined as the percentage of input (1% of lysate) (right); the expression of indicated protein and products of IP was validated by western blot (left). Data are means ± SD analyzed using Student’s *t* test or One-way ANOVA (body weight change); Log-rank test was used to analyze the significance of survival differences. * *p* < 0.05, ** *p* < 0.01, ** *p* < 0.001.

We then used 5Aza for *in vivo* antiviral evaluation in BALB/c mice infected with a mouse-adapted SARS-CoV-2 (MA-SARS-CoV-2) that exhibits high infectivity in mice. In clinical use, the recommended starting dose of 5Aza is 75 mg/m^2^ (equals to ~2mg/kg) in treating MDS. Therefore, mice were intranasally challenged with 2 × 10^3^ plaque-forming units (PFU) of MA-SARS-CoV-2, followed by intraperitoneal administration of 5Aza at 2 mg/kg body weight (equal to 6 mg/m^2^ surface area) at 1 d post infection (dpi), once daily for seven consecutive doses. As shown in Fig.1d, at 5 dpi, mice in 5Aza-treated group started to recovery body weight, while mice body weight in saline-treated group were still decreasing and 5/6 of mice lost more than 25% of body weight. In accordance with this, 83.33% (5/6) of saline-treated animals became moribund (defined as 25% loss of body mass) compared with 16.67% (1/6) in 5Aza-treated group. Besides, the viral RNA copy number and titers in the lungs of 5Aza-treated mice also showed a significant decrease (Fig.1f). Moreover, histological examination revealed remarkable amelioration of lung damage at 4 dpi in the 5Aza group (Fig.S4c). In contrast, the saline group exhibited massive alveolar space mononuclear cell infiltration, moderately severe bronchiolar epithelial cell death, and intra-endothelium and perivascular infiltration of pulmonary blood vessels (Fig.S4b). The RNA-seq of lungs also demonstrated that the 5Aza rescued most downregulated genes with virus infection (Fig.S5), further demonstrated that 2 mg/kg 5Aza treatment protected against SARS-CoV-2 attack. Based on the dose used in mice is equivalent to that in treating MDS in humans, it would be encouraging to extend to human COVID therapy.

As an RNA analog with OH-group on ribose 2’ carbon (2’ C) (Fig.S1a), 5Aza could theoretically incorporate into RNAs. High-resolution mass spectrometry analysis showed that RNA containing 5Aza increased ~40-fold with short-term 5Aza treatment (Fig.S6). We took advantage of 5Aza stability in bisulfite treatment to develop a new method (5Aza-BSseq) that identifies the location where 5Aza is incorporated (Fig.S7). Notably, 5Aza-BSseq results also suggested efficient azacytidine incorporation into viral RNA (Fig.1g).

After deoxidized conversion to decitabine and incorporation into DNA, 5Aza causes endogenous retrovirus (ERV) DNA hypomethylation, which activates retroviral RNA transcription and triggers type I interferon response. In this study, decitabine was five times less efficient than 5Aza in inhibiting viral replication (Fig.S8). Additionally, 5Aza did not significantly increase ERV expression (Fig.S9). Therefore, 5Aza incorporation into RNA might be linked to its inhibitory effects on viral replication.

We further explored the possible antiviral mechanism of 5Aza incorporation into viral RNA. Previous research showed that 5Aza incorporation enhanced lethal mutagenesis on influenza virus. However, we found consistent mutations and comparable mutation rates in the viral RNA propagated in 5Aza-treated or control cells (Supplementary Table 1), excluding the role of 5Aza-induced lethal mutagenesis in anti-SARS-CoV-2. Considering that 5Aza sequesters tRNA methyltransferase to inhibit cytosine-C5 methylation (m^5^C) in tRNA ^1^ (the reaction principle is exhibited in Fig.S10), we further explored whether 5Aza incorporation affects viral RNA methylation. We used nanopore direct RNA sequencing to avoid false-positive methylation sites caused by unconverted 5Aza, as assessed with bisulfite sequencing. The nanopore m^5^C identification algorithm, trained by m^5^C control data as well as many datasets, was used for data analysis. High-confidence m^5^C sites in 5Aza-treated viral RNA decreased significantly by 40% (false positive < 0.05) (Fig.1h). We further validated these m^5^C sites using the optimized RNA-BSseq and strict criteria (Fig.S11 and S12). All m^5^C sites identified through nanopore sequencing were also present in RNA-BSseq (Fig.S13).

The primary writers for m^5^C on mRNAs have been proposed to be NSUN2 and DNMT2, which are demonstrated to contribute to m5C methylation on HIV-1 RNA and thus facilitate virus infection^2,3^. Here, we found that overexpression of DNMT2 and NSUN2, significantly promoted the SARS-CoV-2 replication (Fig.S14), implying both of DNMT2 and NSUN2 are responsible for the m^5^C methylation of SARS-CoV-2 RNA. The immunoprecipitation (IP) assay showed that DNMT2 and NSUN2 could bind to SARS-CoV-2 RNA (Fig. 1i). Notably, DNMT2 IPed more RNAs in the presence of 5Aza. As RNA methylation occurs, cytosine RNA methyltransferases (m^5^C-RMTs) form a covalent thioester bond, connecting the cysteine residue of its catalytic domain to the C6 position, thereby forming an m^5^C-RMT-RNA adduct. Next, RMT transfers a methyl group from the cofactor S-adenosyl methionine (SAM) to C5 of cytosine, followed by enzyme release from the adduct β-elimination. 5Aza is a suicide mechanism-based inhibitor of m^5^C-RMTs^4^. RNAs containing 5Aza at the precise target site will sequester the m^5^C-RMT by generating RNA-m^5^C-RMT adducts, which will result in the decreased level of active endogenous enzymes^5^. Consistent with this theory, we observed an obvious decreased DNMT2, rather than NSUN2 protein upon 5Aza treatment in SARS-CoV-2 infected cells (Fig. S15). It suggests that DNMT2 is more likely the main enzyme for m^5^C RNA methylation in SARS-CoV-2. Further investigations are underway to elucidate the relevance of m^5^C methylation and SARS-CoV-2 infection to 5Aza treatment.

In summary, we demonstrated that 5Aza can incorporate into SARS-CoV-2 RNA and disturb m^5^C RNA methylation modification, potentially contributing to 5Aza’s anti-SARS-CoV-2 activity. We repurposed 5Aza as a promising candidate for combating COVID-19 and introduced the possibility of targeting viral epigenomes as a novel antiviral strategy.

## Acknowledgements

We thank Tao Du, Jin Xiong, and other colleagues from BSL-3 Laboratory of Wuhan Institute of Virology and Wuhan National Biosafety Laboratory for their critical support and excellent coordination. We thank Prof. Meilin Jin of Huazhong Agricultural University for kindly providing mouse-adaptived SARS-CoV-2 strain. We thank Editage (www.editage.cn) for English language editing. This work was supported by the National Science and Technology Major Projects (grant number 2020ZX10001016).

## Competing interest

The authors declare no competing interests.

## Notes

### Competing Interest Statement

The authors have declared no competing interest.

