## supplementary information for "Azacytidine targeting SARS-CoV-2 viral RNA as a potential treatment for COVID-19"

**This Supplementary file includes:**

Materials and Methods

Figures. S1 to S15

**Materials and Methods**

***Cells, virus, and reagents***

Vero E6 cells were obtained from the American Type Culture Collection (ATCC-1586) and cultured in Dulbecco’s modified Eagle’s medium (DMEM; Gibco), supplemented with 10% fetal bovine serum (FBS; Gibco), penicillin (100 U/ml), and streptomycin (100 μg/ml), at 37°C in a humidified 5% CO_2_ incubator.

SARS-CoV-2 (nCoV-2019BetaCoV/Wuhan/WIV04/2019) was propagated in Vero E6 cells. Viral titer was determined through plaque assay. Prof. Meilin Jin from Huazhong Agricultural University provided mouse-adaptive SARS-CoV-2 (MA-SARS-CoV-2), which was created through continuously passaging wild virus in aged mouse lungs^1^. All infection experiments were performed in a biosafety level-3 (BSL-3) laboratory.

5-Azacytidine (5Aza) and Chloroquine (CQ) were obtained from MedChemExpress and dissolved in sterile saline and water, respectively.

Lipo8000 ([Beyotime Biotechnology](http://www.baidu.com/link?url=EC_QjIP4puLIA1A8ux9HkeVOq_9IOsm5YpNLG3xIgPmCaArhnlfFlI3WeR_1ZFsL" \t "_blank), China) was used to transfect plasmids according to the instruction.

***Evaluation of cytotoxicity***

Vero E6 cells were incubated with different 5Aza concentrations in 96-well plates for 24h. Cytotoxicity was measured using CCK-8 assay. EC_50_ was calculated in Prism (GraphPad).

***Quantitative reverse transcription-PCR (qRT-PCR)***

Viral RNA was extracted using the TaKaRa MiniBEST Viral RNA/DNA Extraction Kit (Cat No. 9766). RNA was eluted in 40 μl RNase-free water. Viral RNA copy number was determined with TaqMan-based qRT-PCR using the One Step PrimeScript RT-PCR Kit (Takara, RR064A). To generate a standard curve, the receptor binding domain (RBD) of the spike gene was amplified and its copy number determined from serial dilutions. Primers used for qRT-PCR were RBD-F: CAATGGTTTAACAGGCACAGG, RBD-R: CTCAAGTGTCTGTGGATCACG, and RBD-probe: 5′-FAM-ACAGCATCAGTAGTGTCAGCAATGTCTC-BHQ 3′. The thermocycling schedule was as follows: 95°C for 2 min, followed by 40 cycles of 95°C for 5 s and 60°C for 30 s. The half-maximal inhibitory concentration (IC_50_) of 5Aza was calculated in GraphPad.

***Immunofluorescence microscopy***

Cells were fixed overnight with 4% paraformaldehyde, permeabilized with 0.05% Triton X-10 for 10 min, and blocked with 5% bovine serum albumin (BSA) for 1 h. They were then incubated for 2 h at room temperature with a primary antibody against viral N protein (1:500 dilution, Sino Biological), followed by incubation with Alexa 488-labeled goat anti-rabbit IgG (1:500 dilution, Proteintech) for 1 h. Nuclei were stained with DAPI (Beyotime Biotechnology). Images were captured with fluorescence microscopy.

***Plaque assay***

Vero E6 cells (~1 × 10^5^ cells) were incubated in DMEM containing SARS-CoV-2. After 1 h, the supernatant was removed and the cells were incubated in an overlay medium containing DMEM with 2% FBS and 0.9% CMC (Calbiochem). After 96 h, the cells were fixed overnight using 4% paraformaldehyde and stained with crystal violet.

***Time-of-drug-addition assay***

A time-of-drug-addition assay was performed to determine the SARS-CoV-2 life-cycle stage that 5Aza affects. For “Entry,” Vero E6 cells were pretreated with 32 μM 5Aza for 1 h and infected with 0.2 MOI SARS-CoV-2. Two hours later, the cells were washed twice with DMEM and placed in fresh culture medium. For “Post-entry,” the cells were infected with virus for 2 h, then washed twice and cultured with fresh medium containing 32 μM 5Aza. For “Full-time,” cells were pretreated with 32 μM 5Aza for 1 h and infected with SARS-CoV-2 for 2 h. Subsequently, the cells were washed twice and cultured with fresh medium containing 32 μM 5Aza until the experiment ended. Next, supernatants were collected, cells were fixed, and total proteins were extracted for qRT-PCR, fluorescence microscopy, and western blotting, respectively.

***RNA immunoprecipitation***

Vero E6 cells seed in 10-cm plate were transfected with pCAGGS/HA expressing eGFP, DNMT2, or NSUN2. 24 hours later, cells were infected with 0.2 SARS-CoV-2 for 20 hours in the presence of 16 μM 5Aza or not. Cells were lysed with RIP lysis buffer (gzscbio, Guangzhou, China) with inhibitor cocktail. 1% of the lysates was collected for input analysis, and 3% of lysates were collected for WB analysis. RNAs of remaining lysates were immunopricipated by anti-HA agarose in the presence of RNase inhibitor and 0.2 M EDTA at 4°C overnight. After washing with RIP buffer for 8 times, 20% of the agarose was denatured for IPed protein detection, the remaining was subjected to 10% SDS and proteinase K treatment at 65°C for 45 min. Then the supernatant was collected for RNA extraction, followed by RT-PCR for SARS-CoV-2 RNA detection.

***Western blotting***

Cells were lysed with RIPA buffer (Thermo, 89900) containing a protease inhibitor cocktail (MCE). Proteins were separated using 10% SDS-PAGE and then transferred onto PVDF membranes (Millipore). Membranes were blocked in TBS buffer with 0.1% Tween-20 (TBST) for 1 h at room temperature, followed by incubation with a primary antibody against viral N protein for 2 h at room temperature. After washing with TBST three times, the membranes were incubated with a secondary peroxidase-conjugated antibody (1:5000 dilution, Proteintech). GAPDH was the internal control. Bands were detected using WesternBright ECL (Advansta).

***Animal experiments***

Female BALB/c mice (6–7 weeks old) were obtained from Snac Jinda Laboratory Science (Hunan Province, China) and randomly divided into three groups (mock infection, SARS-CoV-2+saline, and SARS-CoV-2+5Aza; n = 9 per group). On Day 0, mice were anesthetized using isoflurane. Infection groups were intranasally challenged with 2 × 10^3^ PFU MA-SARS-CoV-2 in 50 μl DMEM, while the mock infection group was treated with 50 μl DMEM only. One day later, mice were intraperitoneally injected with 2 mg/kg 5Aza (SARS-CoV-2+5Aza group) or an equivalent volume of sterile saline (SARS-CoV-2+saline and mock infection groups), once daily for seven consecutive doses. Body weight was measured daily for 7 d (n=6); mice with more than 25% of body weight mass were considered dead, and mouse survival rates were calculated (n=6). At 4 dpi, three mice from each group were euthanized for lung collecting. Tissues were used for the qRT-PCR quantification of viral RNA copy number, plaque assay of virus titer, and H&E staining for histopathological changes.

All animals were bred in a specific pathogen-free animal facility; experiments were performed in a biosafety level-3 (BSL-3) laboratory and were approved by the Institutional Animal Care and Use Committee of Wuhan Institution of Virology, CAS.

***Bisulfite library preparation, RNA library preparation and next-generation sequencing***

Total RNA (200 ng) was treated with the MGIEasy rRNA Depletion Kit for rRNA removal, bisulfite-converted using the EZ RNA Methylation Kit (Zymo Research), then fractionated into long and small RNA fractions.

The transcriptome library of bisulfate RNA or non-bisulfate RNA was constructed using the MGIEasy RNA Directional Library Prep Kit (without ligase in the second strand synthesis). Samples were sequenced using the PE100+10 bp method. ERCC was used as negative control and helped to correct false positive sites. The ERCC without any methylation could be used to assay the RNA bisulfite conversation rate.

***Nanopore direct RNA sequencing***

Total RNA (1 µg) was treated with the MGIEasy rRNA Depletion Kit to remove rRNA. Next, a direct RNA Sequencing kit (SQKRNA002) was used to prepare RNA with 1D sequencing on an Oxford Nanopore equipment. Data were analyzed using a pipeline established in the Zhang lab.

***Quality control***

First, a custom Python script was used to demultiplex the barcode list. Second, adapter trimming and quality pruning were analyzed with fastp. Bases in the sliding window with mean quality below 20 were cut, and raw reads of those containing 5 or 3 bp adapters were trimmed. mRNA was trimmed to nine bp from the left end.

***Mapping and calling***

C2T and G2A reference genomes were converted for building indexes using bismark _genome_preparation. Clean sequencing reads were aligned to the two reference genomes with Bismark. Methylation calls were stored in BAM files generated during this step. BAM files were used to ensure that reads mapped to the same direction and had the same start positions, mapping length, and molecular tag. Bismark_methylation_extractor was used to obtain readable results for downstream analysis.

***Quantity and differential expression***

Transcript quantification was performed using HTSeq v0.44.0. RNA with zero counts in all samples was removed. Normalization and differential expression were determined using the R package DEseq2 v1.28.1.

***Differentially methylated regions***

After alignment, we extracted methylation counts from Bismark output; then, DMRfinder was used to cluster CpG sites into test regions for differential methylation.

***Sample preparation for nucleotide modification***

For DNA and RNA extraction, cells were lysed with TRIzol reagent and digested with proteinase K at 37°C for 30 min. An equal volume of back extraction buffer (4 M guanidine thiocyanate, 50 mM sodium citrate, 1 M Tris, pH 8.0) and 1/10 volume of 3 M sodium acetate (pH 4.0) was mixed with the cell lysate. Samples were then centrifuged at 13,200 rpm for 15 min at 4°C. The upper phase was transferred to a new tube and an equal volume of 100% isopropanol was added. After overnight incubation at -80°C, samples were centrifuged at 13,200 rpm and 4°C for 15 min. The supernatant was removed and pellets were washed twice with 70% ethanol. Samples were eluted with 50 μl nuclease-free buffer and centrifuged at 5,000 rpm for 2 min. The supernatant was then transferred to a new tube. Total RNA was extracted using TRIzol reagent. RNA or DNA/RNA hybrid (500 ng) was digested into nucleotides using 20 μl Nucleoside Digestion Mix (NEB, M0649S). The digestion product was brought to 50 μl with nuclease-free water and filtered using a Microcon-10kDa Centrifugal Filter Unit with Ultracel-10 membrane.

***LC-MS/MS***

The analysis used an ACQUITY UPLC I-Class system coupled with a QTRAP 6500+ triple quadrupole mass spectrometer (Sciex, Framingham, MA). Nucleotide samples were diluted five times, and 10 μl of each dilution was chromatographed on HSS T3 C18 (100 x 2.1 mm, 1.8 μm, Waters, Milford, MA). Samples were eluted with a linear gradient of A (0.2% formic acid + 5 mM ammonium formate in deionized water) and B (0.2% formic acid + 5 mM ammonium formate in methanol) at a flow rate of 0.3 ml/min and temperature of 45°C. The gradient elution started with 96% of mobile phase A for 2.5 min. Next, mobile phase B was linearly increased up to 45% in 2.5 min and maintained for 0.5 min, then linearly increased up to 100% in 2 min. Finally, the combination was brought back to 96% A in 1 min. Detection was achieved using a QTRAP 6500+ system in positive ion mode and multiple reaction monitoring mode (MRM). Source parameters were set as follows: temperature, 400°C; curtain gas (CUR), 30; collision gas, low; ion spray voltage, 5000 V; ion source gas (GS1), 45; and drying gas (GS2), 40.

***Statistical analysis***

Statistical significance was determined by Student’s *t* test for two groups. Analysis with a probability value < 0.05 were considered statistically significant.

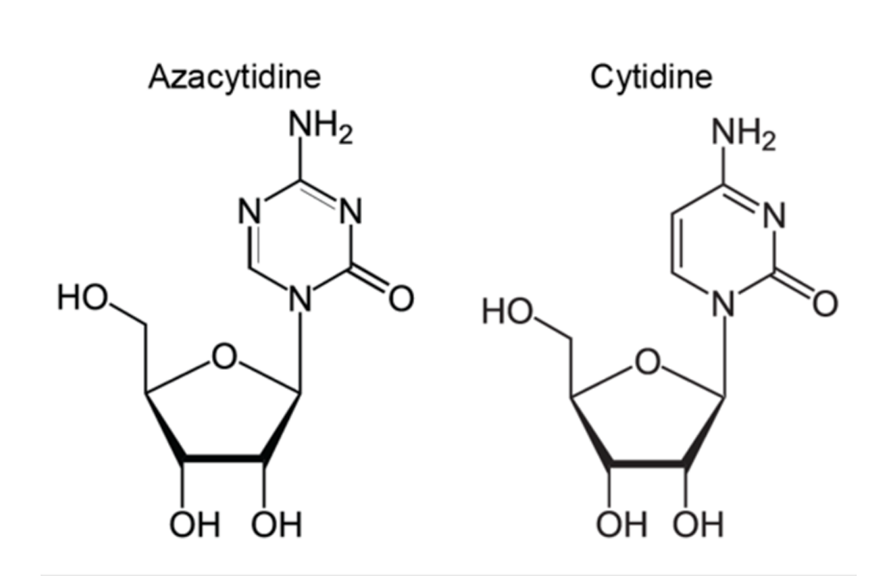
**Supplementary figures**

**Fig.S1 Molecular structures of cytidine and azacytidine.**

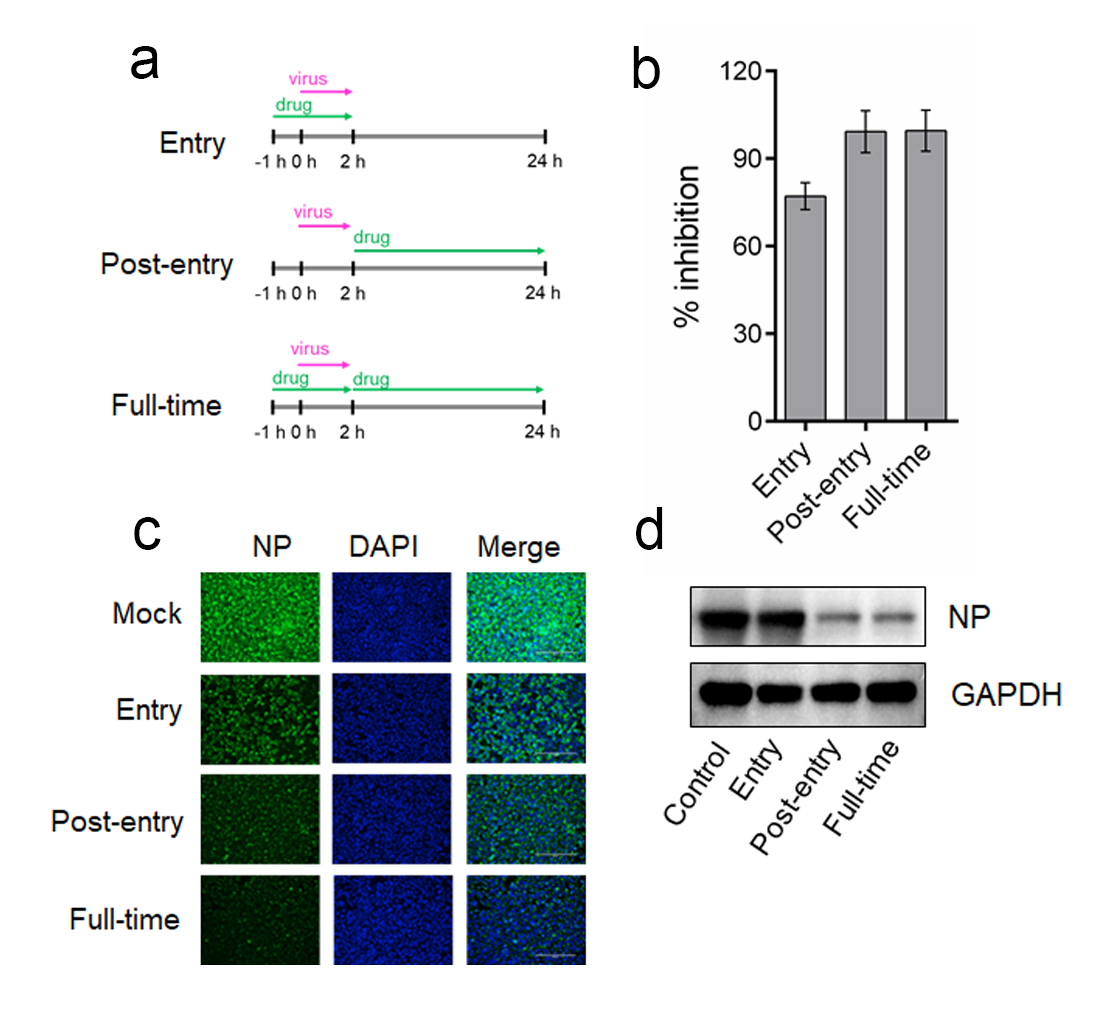
**Fig.S2 Time-of-drug-addition experiment of 5Aza.**

**a** Schematic of time-of-drug-function experiment of 5Aza. Vero E6 cells were treated with 5Aza and then infected with 0.2 MOI SARS-CoV-2. **b** After 24 hpi, supernatant was collected for detecting RNA copies using qRT-PCR. **c** cells were fixed for immunofluorescence assay of viral N protein, and **d** total proteins were extracted for western blotting to detect N protein. Nuclei were stained with DAPI. Bars, 200 μM. All experiments were performed in triplicate; data are mean ± SD, and analyzed using Student’s *t* test.

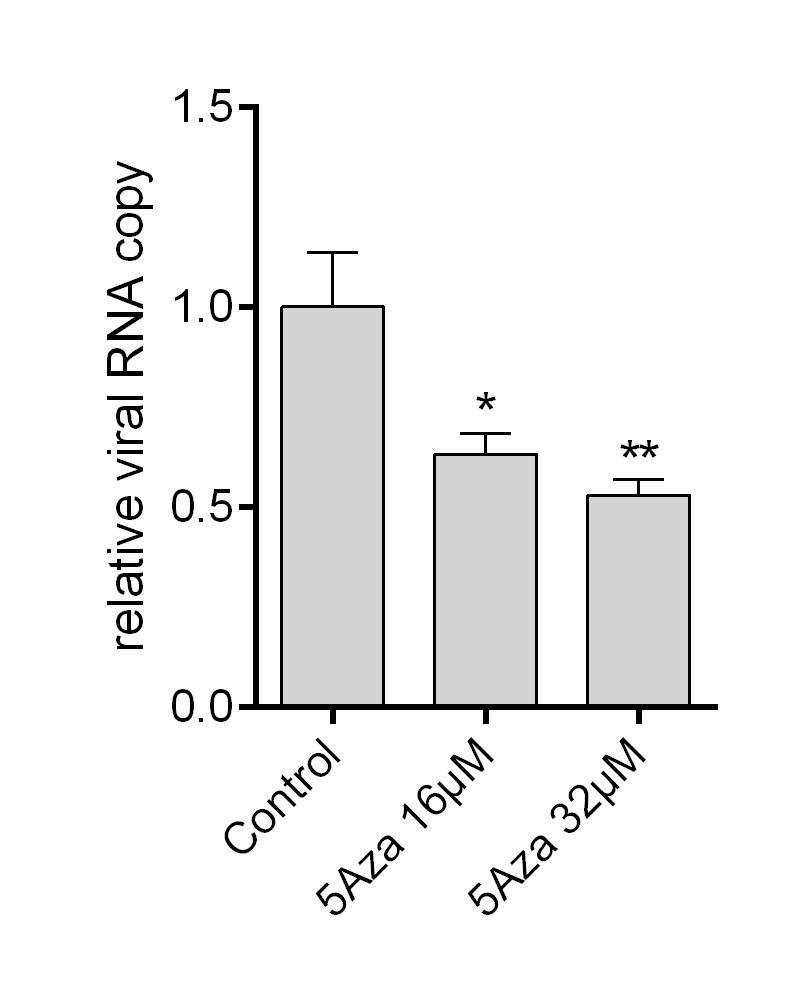
**Fig.S3 The anti-SARS-CoV-2 effect of** **5Aza when the drug was added 24 hours after virus infection.**

Vero E6 cells were infected with 0.1 moi SARS-CoV-2, 24 hours later, various concentration of 5Aza was added. Another 24 h later, the supernatant was collected for viral RNA detection by qRT-PCR. Data are mean ± SD, and analyzed using Student’s *t* test.

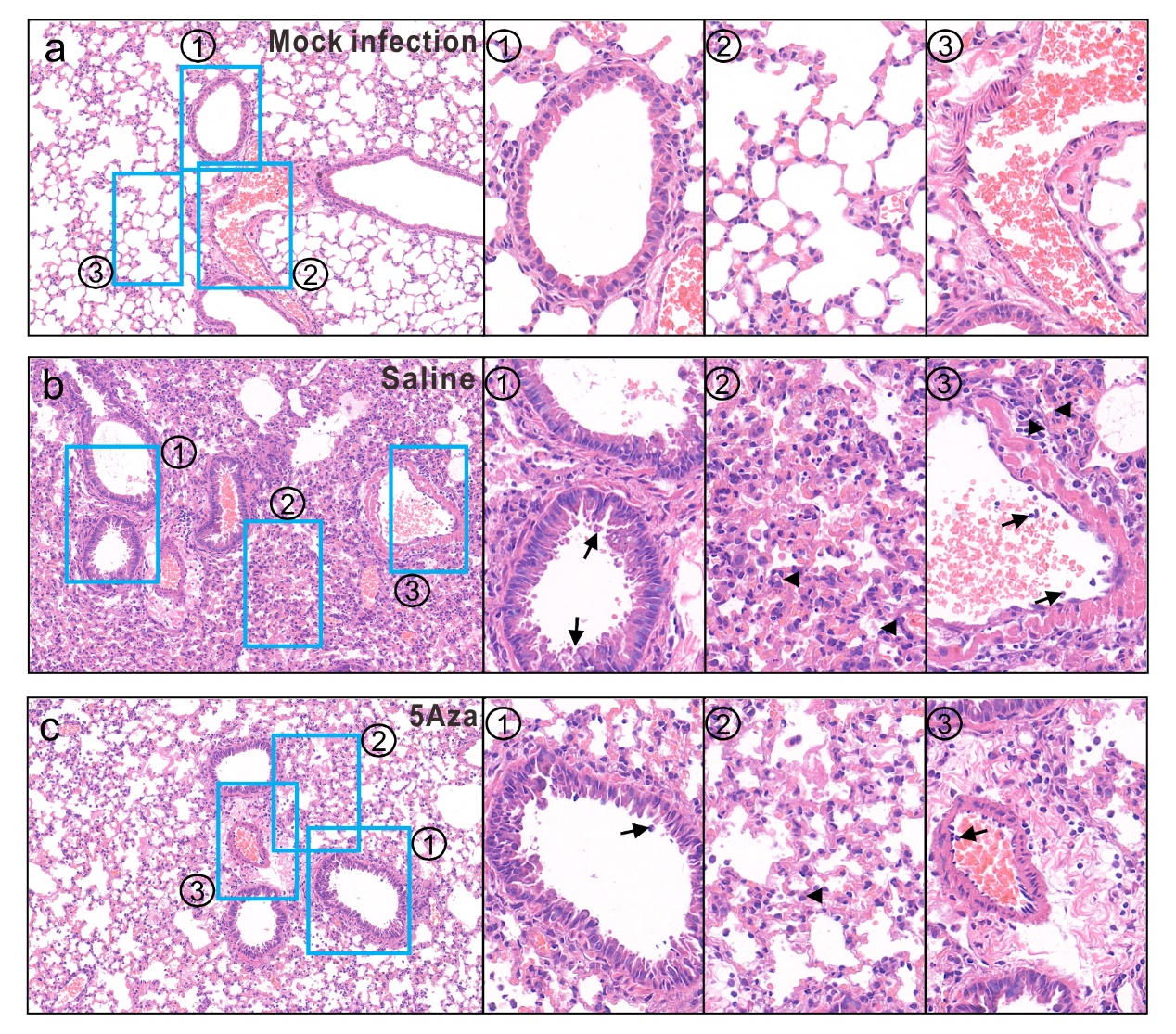
**Fig.S4 H&E staining of lung tissues.**

At 4 dpi, mice were euthanized, and lungs were collected for histopathological analysis with H&E staining. Numbered blue rectangles are shown in magnified images to the right. **a** Lung tissues were normal in the mock infection group. **b** Lung tissue section from the saline group mice displayed (1) bronchiolar epithelium cell death (arrow), (2) destruction of alveola with massive infiltration (arrowhead), (3) endothelium infiltration in blood vessels (arrows) and perivascular infiltration (arrowhead). **c** Representative image of lung tissue from 5Aza mice showing alleviated infiltration.

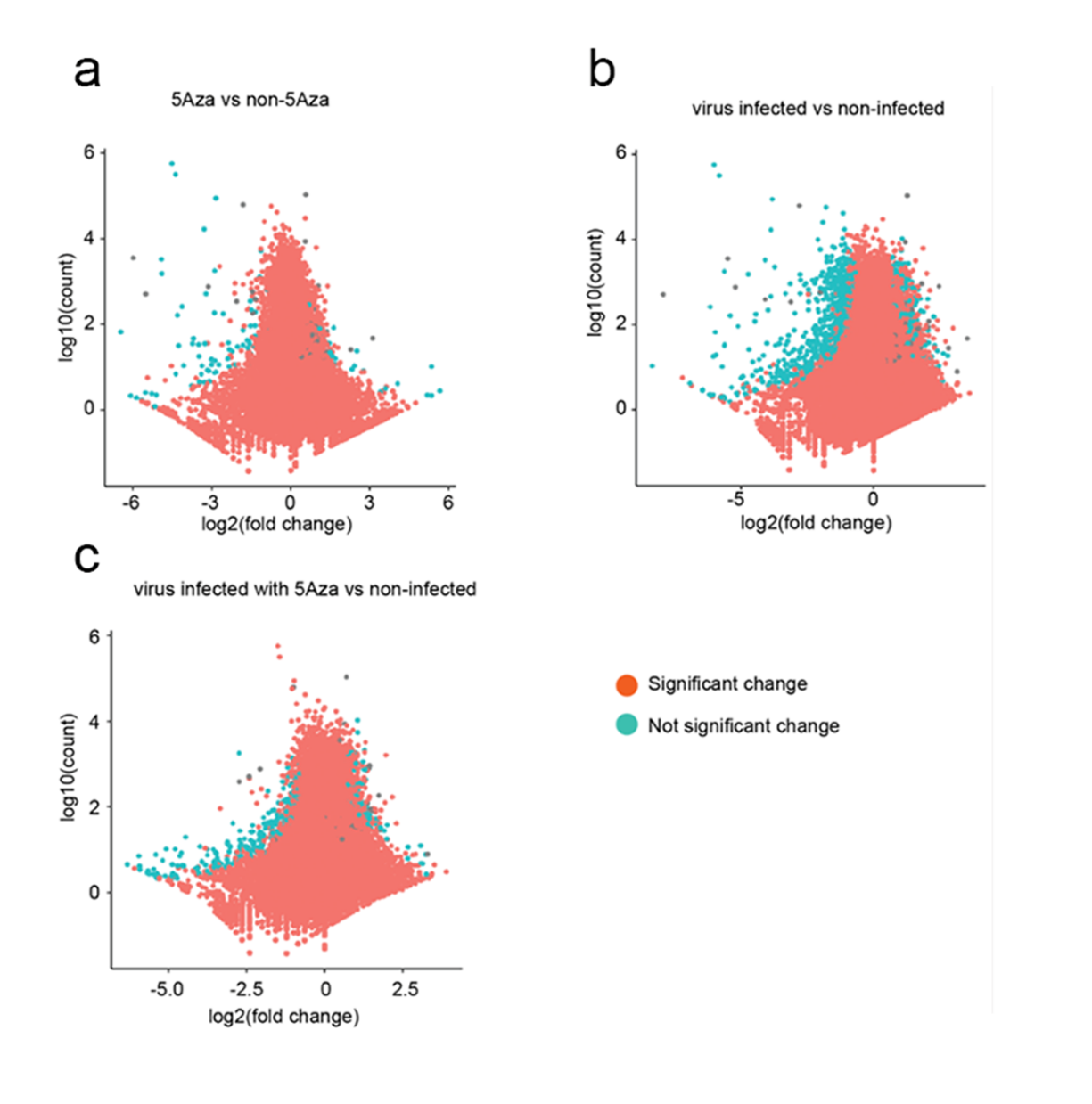

**Fig.S5 RNA-seq for the lung tissue of the mice.**

**a** The mice were treated with 5Aza (2mg/kg) and saline buffer for 4 days. The lung tissues were collected and RNA was extracted for RNA-seq. The x-axis showed the log2(5Aza treated sample/control sample). The y-axis indicated log10(mean counts on genes). The volcano plot showed that the 5Aza did not change the gene expressions significantly. **b** Mice were challenged by MA-SARS-CoV-2 and non-infected. The lung tissues were collected at 4dpi and RNA was extracted for RNA-seq. The x-axis showed the log2(virus infected sample/non-infected sample). Most of the genes were downregulated significantly after virus infection. **c** Mice were challenged with MA-SARS-CoV-2 then treated by Aza (2mg/kg) on 1dpi. The lung tissues were collected at 4dpi and RNA was extracted for RNA-seq. The x-axis showed the log2(Aza treated virus infected sample/non-infected non-5Aza-treated sample). Most downregulated genes in **b** were rescued by 5Aza treatment.

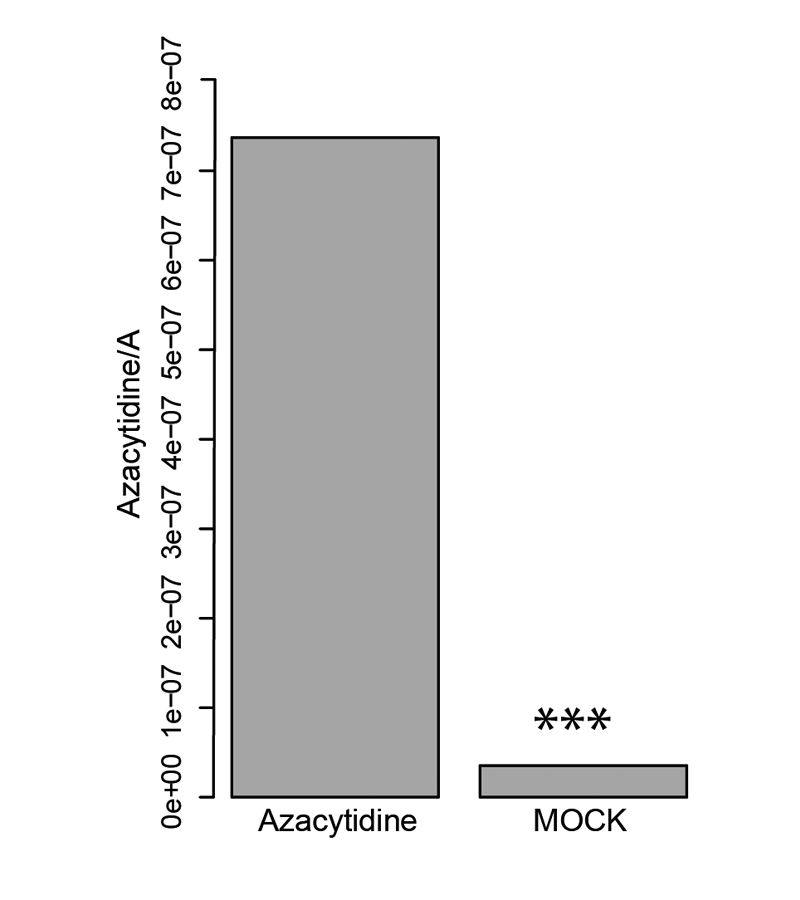
**Fig.S6 Profiling and quantification of azacytidine in DNA and RNA using liquid chromatography-tandem mass spectrometry (LC-MS/MS).**

Considering azacytidine as an RNA analog with OH-group on ribose 2' carbon (2' C), it could theoretically incorporate into RNAs. “Azacytidine” sample were the cells incubated with 5Aza (10 μM) for 24 h and “Mock” sample were the cells without any treatment. Both DNA and RNA were extracted, followed by single nucleotide digestion (DNA and RNA) and LC-MS/MS. Azacytidine was normalized to adenosine on the y-axis. Data are presented as means of triplicate assays (****p* < 0.001). Most of the azacytidine incorporated into the RNAs. This was consistent with previous reports, in which radiolabelled azacytidine was measured by scintillation-based quantification or high-resolution mass spectrum termed "AZA-MS"^2^.

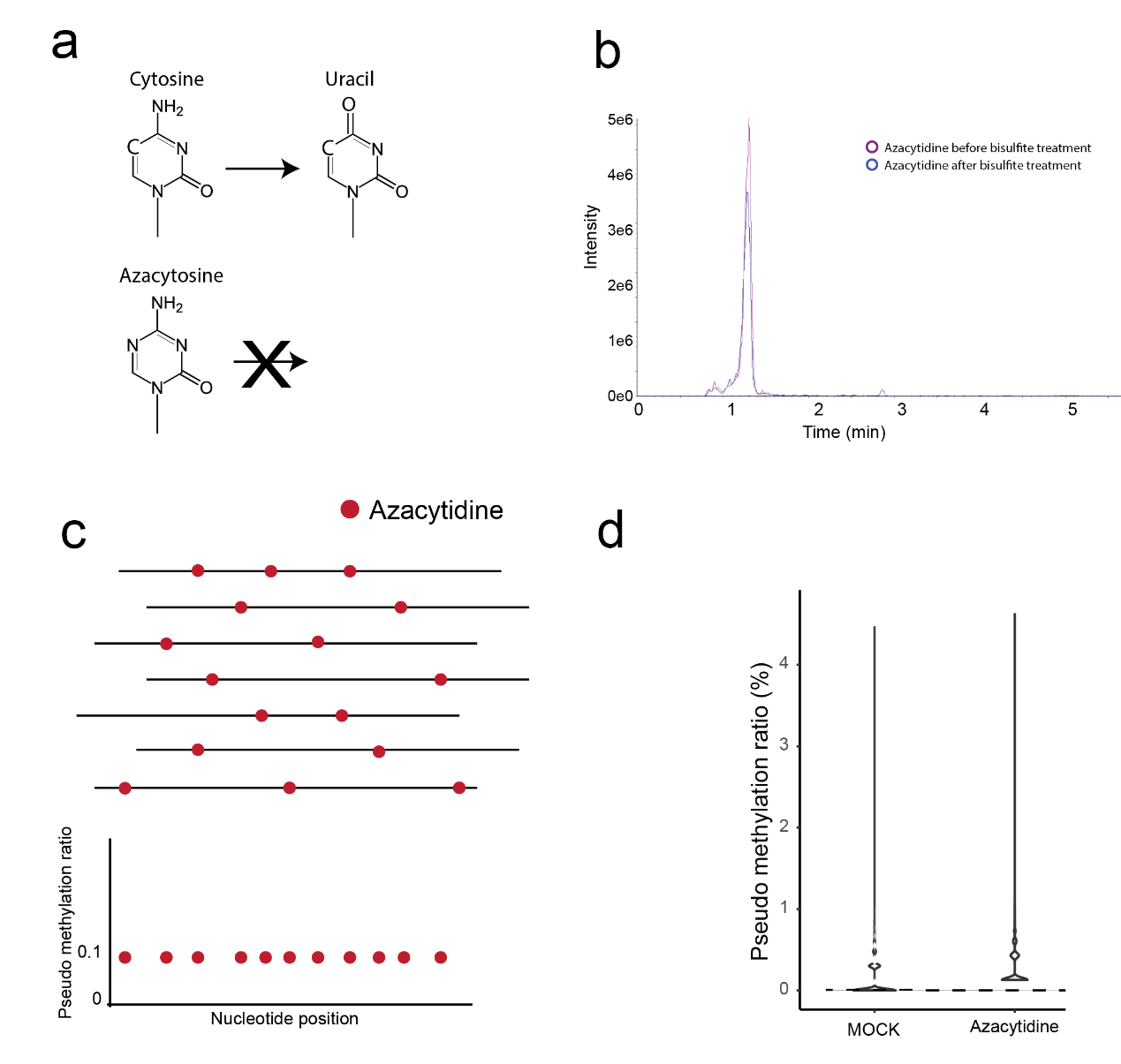
**Fig.S7** **5Aza-BSseq-identified locations of 5Aza incorporation.**

**a** We explored whether azacytidine could be converted to uracil under bisulfide treatment and found that was not the case. Bisulfite deamination of cytidine relies on Lewis acid formation and electron transfer from C5. In azacytidine, N5 inhibited this deamination. b LC-MS chromatograms of 5Aza before and after bisulfite treatment. Supporting the observation in **a**, LC-MS chromatograms of azacytidine remained unchanged before and after bisulfite treatment (). **c** Schematic diagram of testing pseudo-methylation sites. This chemical property of azacytidine is similar to the RNA m^5^C, which also cannot be converted to uracil in bisulfite sequencing. Unconverted azacytidine may be identified as a pseudo cytidine (pseudo-methylation) site in RNA bisulfite sequencing. Owing to randomized azacytidine incorporation, these azacytidine (pseudo-methylation) sites should be observed as increased ultra-low methylation (methylation ratio <1%). **d** The ratio of pseudo-methylation. Pseudo-methylation distribution analysis revealed that azacytidine treatment increased ultra-low pseudo-methylation (<1%), indicating 5Aza incorporation in RNA.

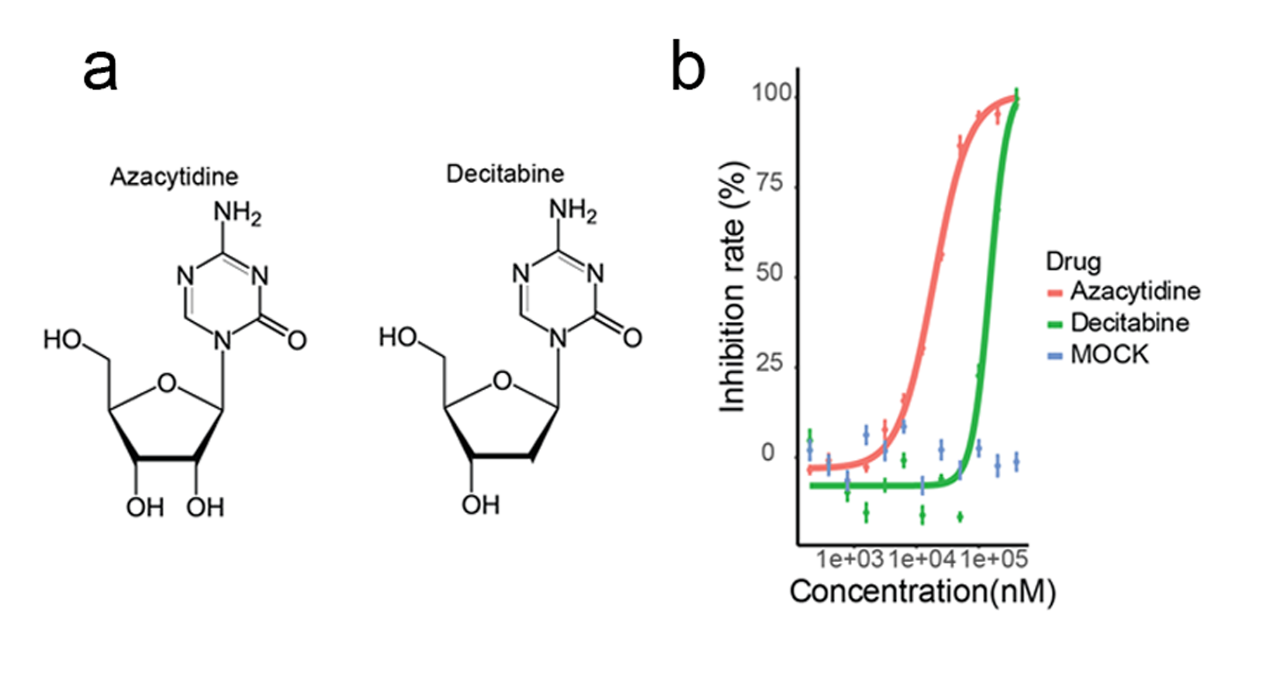
**Fig.S8 Effects of azacytidine and decitabine on SARS-CoV-2 infection.**

**a** Decitabine (5-aza-2'-deoxycytidine) is the deoxidized form of azacytidine. In theory, most decitabine incorporated into DNA and most azacytidine incorporated into RNA. Some azacytidine could be converted to 5-aza-2'-deoxycytidine (decitabine) through oxidization pathway and incorporated into DNAs ^3^. Therefore, the azacytidine antivirus activity could be explained by two possible mechanisms: DNA incorporation or RNA incorporation. **b** Azacytidine and decitabine inhibited SARS-CoV-2 virus infection. Vero E6 cells were infected with SARS-CoV-2 (MOI = 0.2), at different drug concentrations, for 24 h. Supernatant was collected for qRT-PCR-based detection of viral RNA copy number, which allowed for assessment of SARS-CoV-2 inhibition rate. Decitabine has the 5x lower antivirus activity than the azacytidine, suggesting that the azacytidine antivirus activity may rely on RNA incorporation. Data are means ± SD. Solid lines are nonlinear regression curves created in Graphpad.

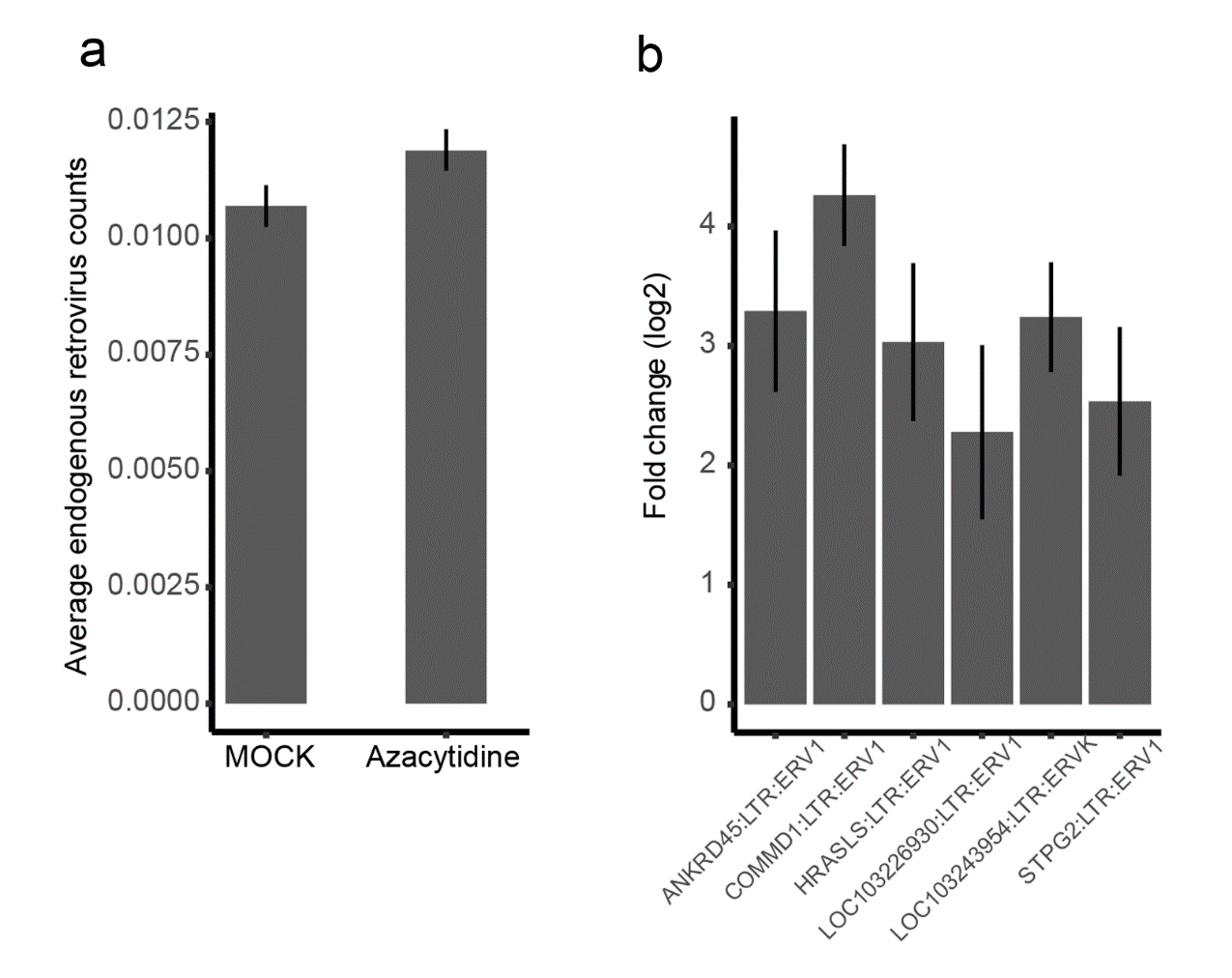
**Fig.S9** **The endogenous retrovirus (ERV) gene expression with ot without 5Aza.**

The previous studies also demonstrated that, after conversion to decitabine and incorporation into DNA^4^, azacytidine causes endogenous retrovirus (ERV) DNA hypomethylation, which activates retroviral RNA transcription and triggers the type I interferon response and viral defense pathway in host cells^5^. Therefore, we further explored the endogenous retrovirus gene expression. Vero E6 cells infected with 0.2 moi SARS-CoV-2 were treated with or without 10 μM 5Aza. After 24 h, cellular total RNA was extracted. **a** Expression of ERV genes in the control and 5Aza-treated samples, determined via RNA sequencing. Average ERV counts did not differ significantly between the control and azacytidine samples. **b** Significant fold change (log2) in differentially expressed ERV genes between the azacytidine and control samples. Only six endogenous retrovirus genes are significantly upregulated. Fold changes were calculated in Deseq2. 5Aza treatment did not increase endogenous retrovirus (ERV) gene expression.

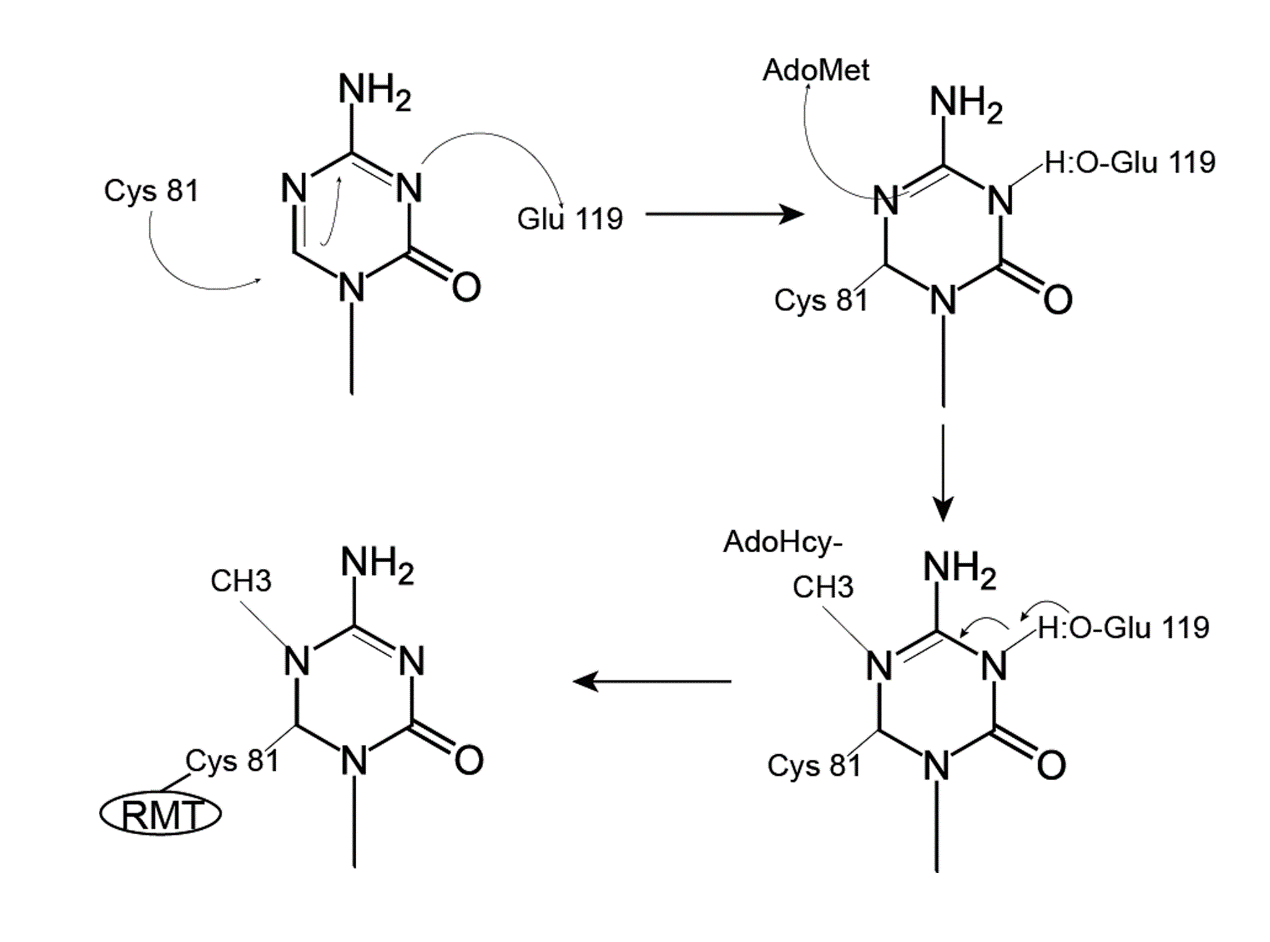
**Fig.S10** **RNA methyltransferases (RMTs) catalyze the methylation of ^5^N in azacytosine.**

RMT first forms a covalent thioester bond to connect the cysteine residue of its catalytic domain to the ^6^C position of azacytosine and then transfers a methyl group from cofactor SAM to ^5^N of azacytosine. Due to ^5^N methylation, enzyme release cannot occur through β-elimination. Therefore, azacytidine can irreversibly sequester RMTs^6^.

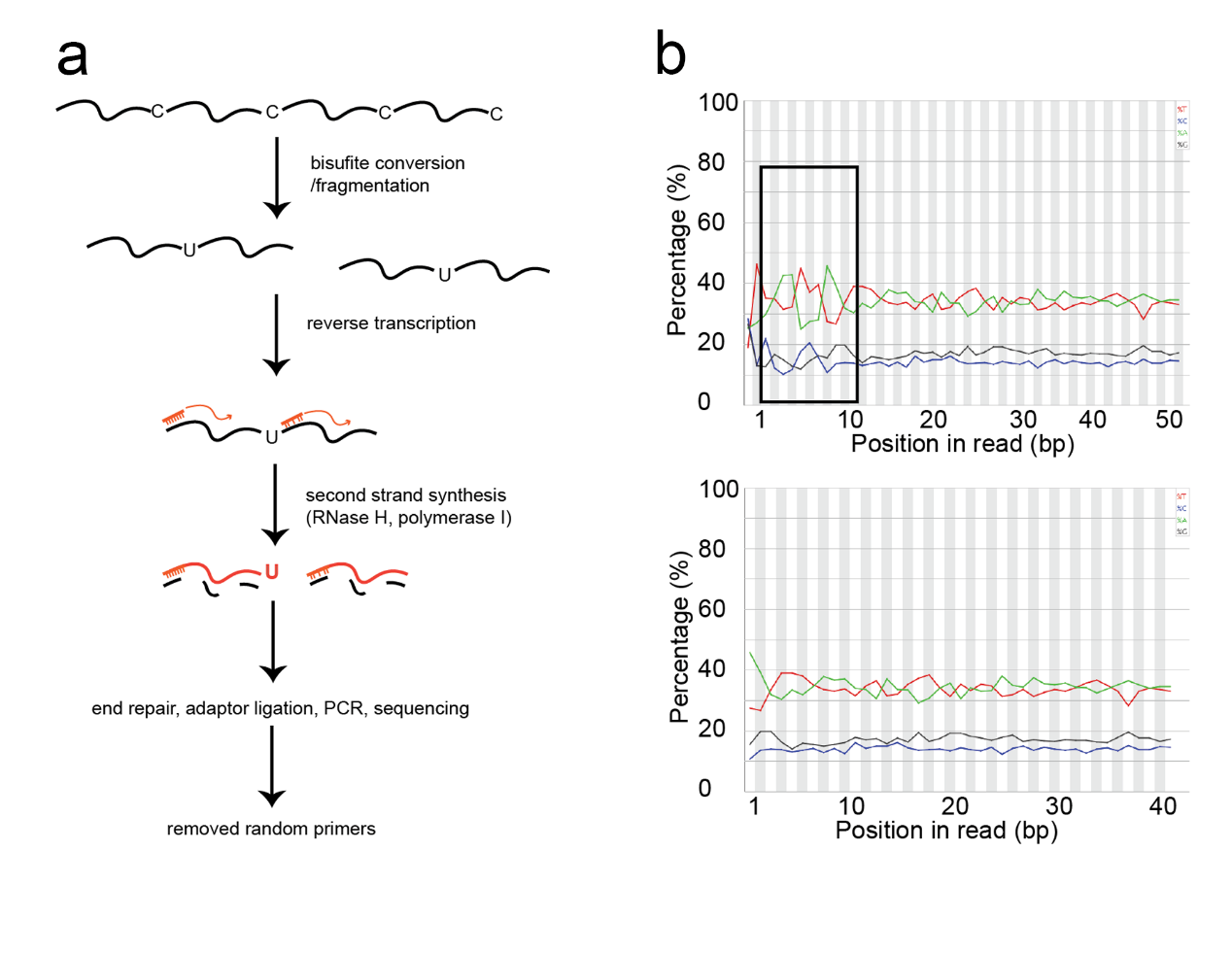
**Fig.S11 Experimental flowchart of RNA-BSseq.**

Limitations in site-mapping technology hamper the validity of coronavirus RNA methylation. As random priming in RNA bisulfite sequencing introduces mismatch errors, it is difficult to reliably measure m^5^C sites. We sought to develop an optimized approach that avoids random primer insertion into sequenced fragments, thus minimizing false positives. (a) Without using ligase in second strand synthesis, we could remove random primers in the middle of sequenced fragments. During bioinformatics analysis, we could then easily detect and delete the first six bases affected by random primers initializing reverse transcription. (b) Nucleotide density distribution (A, T, C, or G) in sequencing reads. Upper and lower panels show distribution before and after the removal of random primer sequences, respectively. The distribution fluctuated before and after a smooth line, indicating a few artifact Cs in our modified bisulfite sequencing.

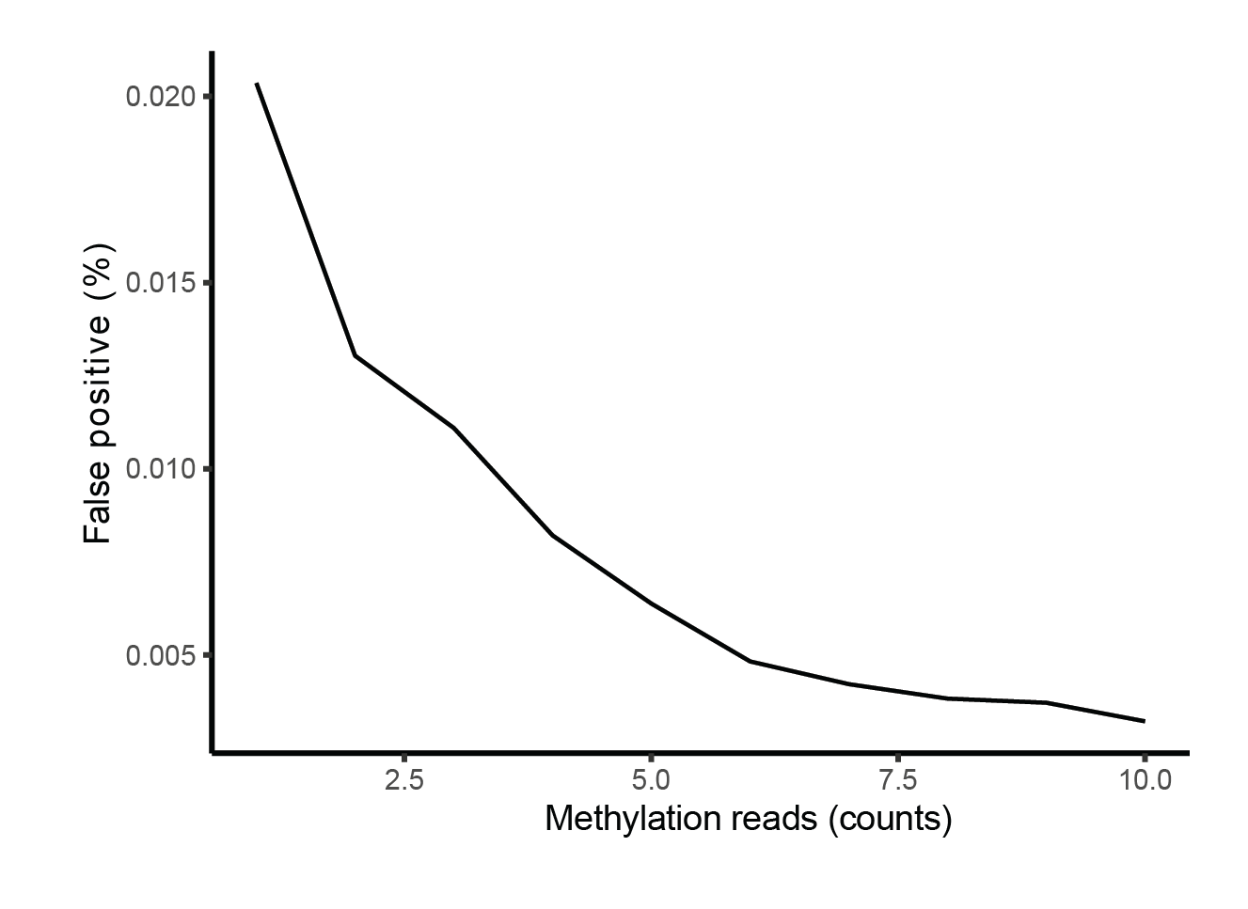
**Fig.S12** **False-positive rate of ERCC (External RNA Controls Consortium) RNA bisulfite sequencing.**

ERCC RNA was synthesized from RNA without m^5^C, subjected to bisulfite treatment, and used for library construction. To further enhance the accuracy of RNA-BSseq detection, we used the ERCC to test the false positive rate of our method and to select a suitable cut-off (coverage > 5, methylation ratio > 95%, false positive~0.005).

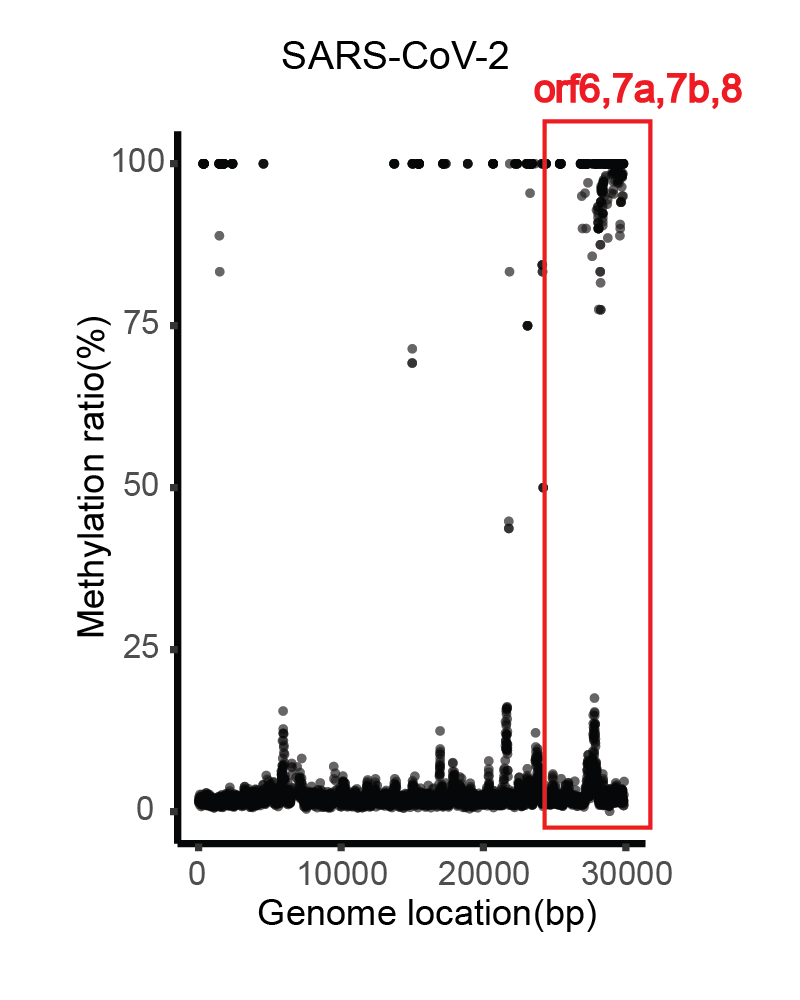
**Fig.S13 RNA-BSseq identified m^5^C locations on SARS-CoV-2 genome.**

There were five minimum methylation supporting reads per point. We found over 555 high-confidence m^5^C methylation sites (methylation ratio > 95%) (false positive = 0.005, bisulfite conversion rate = 97.37%). The red square indicates high methylation in the SARS-CoV-2 genome regions orf6, 7a, 7b, and 8. Data are averaged from three technological repeats.

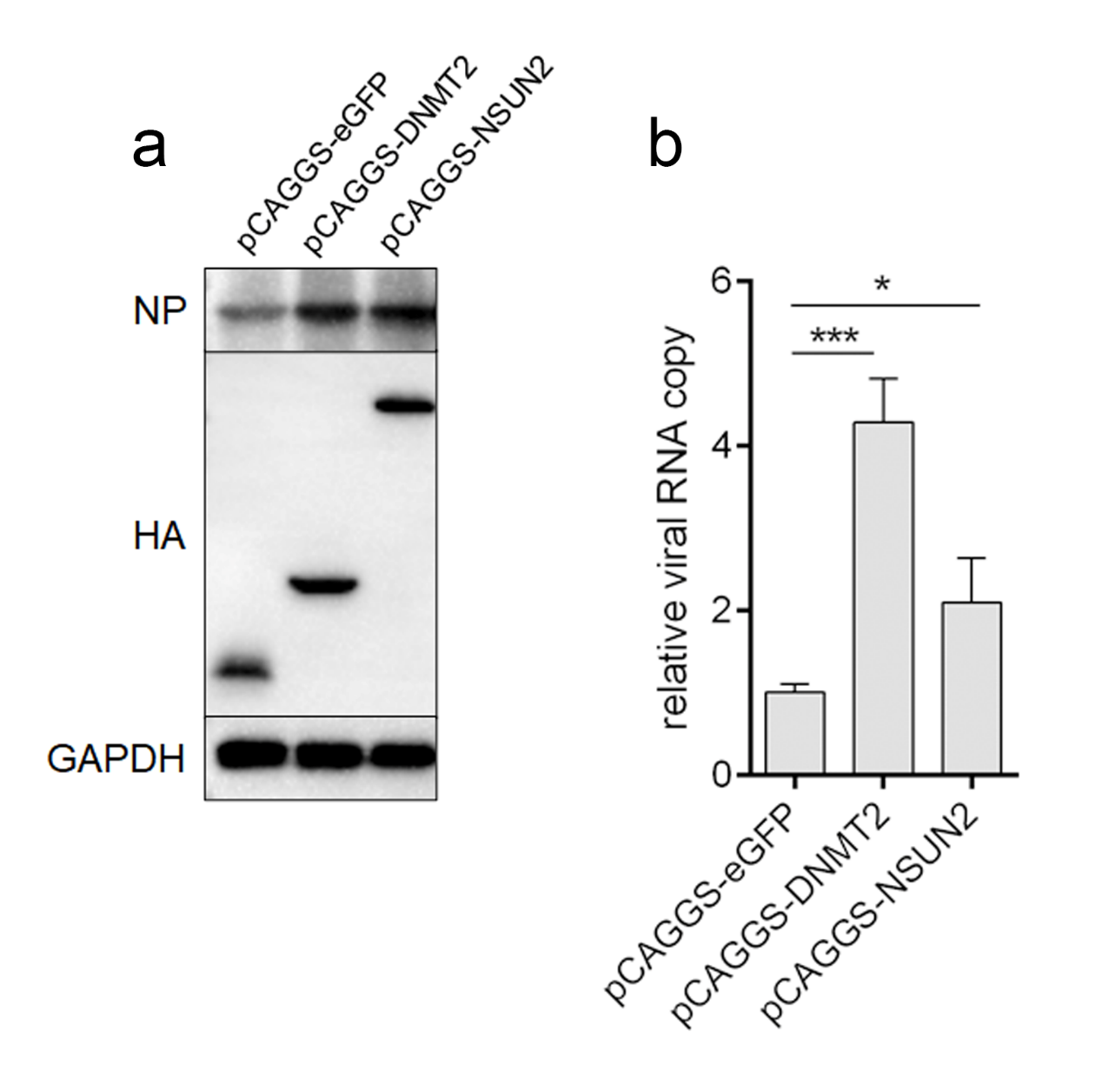

**Fig.S14 overexpression of DNMT2 and NSUN2 promoted SARS-CoV-2 infection.**

Vero E6 cells were transfected with pCAGGS/HA-eGFP, DNMT2, or NSUN2. 24 hours later, cells were infected with 0.1 moi SARS-CoV-2. After 24 h, the supernatant was collected for viral RNA detection by RT-PCR. **b** Cells were lysed with RIPA buffer, and lysates were subjected to western blot for detecting indicated proteins. Experiment was performed in triplicate; data are mean ± SD, and analyzed using Student’s *t* test.

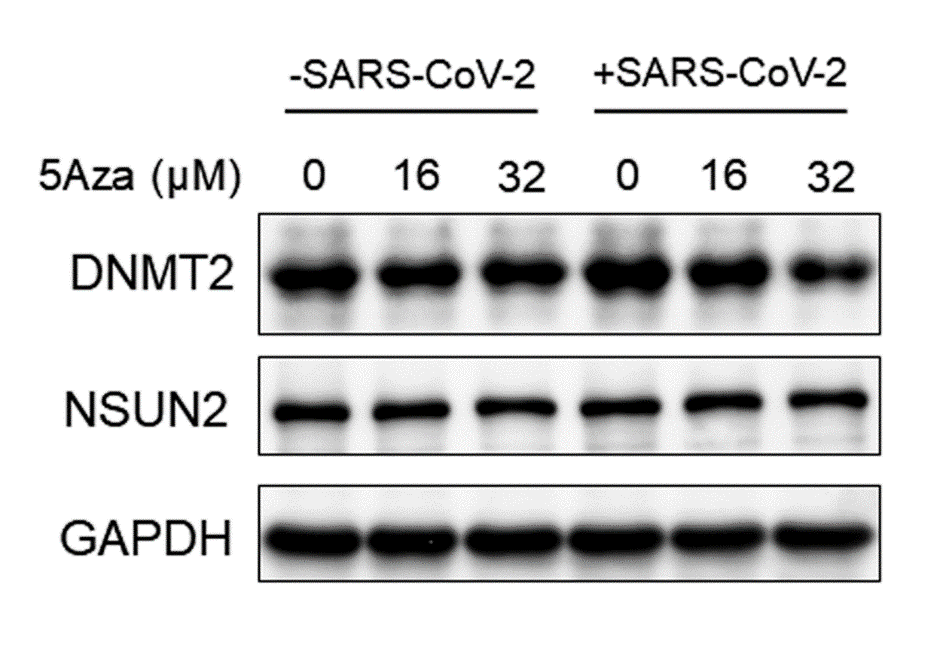

**Fig.S15 The effect of 5Aza treatment on the expression of DNMT2 and NSUN2 protein.**

Mock or SARS-CoV-2 (moi=0.1) infected Vero E6 cells were treated with indicated concentration of 5Aza. After 24 h, cells were lysed with RIPA buffer, and the lysates were subjected to western blot for detecting indicated proteins.
